# Amygdalar and ERC Rostral Atrophy and Tau Pathology Reconstruction in Preclinical Alzheimer’s Disease

**DOI:** 10.1101/2022.06.06.494859

**Authors:** Kaitlin M. Stouffer, Claire Chen, Sue Kulason, Eileen Xu, Menno P. Witter, Can Ceritoglu, Marilyn S. Albert, Susumu Mori, Juan Troncoso, Daniel J. Tward, Michael I. Miller

## Abstract

Previous research has emphasized the unique impact of Alzheimer’s Disease (AD) pathology on the medial temporal lobe (MTL), a reflection that tau pathology is particularly striking in the entorhinal and transentorhinal cortex (ERC, TEC) early in the course of disease. However, other brain regions are affected by AD pathology during its early phases. Here, we use longitudinal diffeomorphometry to measure the atrophy rate from MRI of the amygdala compared with that in the ERC and TEC in controls, individuals who progressed from normal cognition to mild cognitive impairment (MCI), and individuals with MCI who progressed to AD dementia, using a dataset from the Alzheimer’s Disease Neuroimaging Initiative (ADNI). Our results show significantly higher atrophy rates of the amygdala in both preclinical and MCI ‘converters’ compared to controls, with rates of volume loss comparable to rates of thickness loss in the ERC and TEC. Using our recently developed method, referred to as Projective LDDMM, we map measures of neurofibrillary tau tangles (NFTs) from digital pathology to MRI atlases and reconstruct, for the first time, dense 3D spatial distributions of NFT density within regions of the MTL. The distribution of NFTs is consistent with the MR atrophy rates, revealing high densities not only in ERC, but in the amygdala and rostral third of the MTL. The similarity of the location of NFTs and shape changes in a well-defined clinical population suggests that amygdalar atrophy rate, as measured through MRI may be a viable biomarker for AD.

**PACS:** 0000, 1111

**2000 MSC:** 0000, 1111

**Highlights:**

- Amygdala atrophy rate estimated from MRIs in preclinical Alzheimer’s disease (AD)
- 3D distributions of neurofibrillary tau tangles (NFTs) reconstructed from 2D histology
- NFTs highest in the rostral medial temporal lobe, including amygdala and ERC
- Amygdala atrophy rate is comparable to ERC atrophy rate in preclinical AD
- Amygdala and ERC atrophy rates as potential biomarkers rooted in NFT pathology

## 1. Introduction

Alzheimer’s disease (AD) is the leading cause of dementia worldwide [1]. Diagnosis and characterization of AD in its *early* stages remain key challenges, as existing technologies limit the identification of the neuropathological patterns thought to emerge years before symptom onset [2, 3, 4]. In clinical practice, AD is typically first characterized by progressive clinical changes in memory and behavior, and subsequently through imaging changes that indirectly reflect AD neuropathology (i.e. misfolded proteins, tau and amyloid-Beta (A*β*)) [5, 6, 7]. Efforts to identify and understand the spatiotemporal profile of AD in its early stages have centered on these biomarkers [8]–measures that indirectly reflect the underlying pathology, which are obtainable over the course of disease. Of the methods used, neuroimaging has emerged as a prominent player with the ability to localize pathology noninvasively (e.g. tau/amyloid PET) [9, 10], and with proposed surrogates such as shape diffeomorphometric markers (e.g. MRI) [11, 12, 13]. While these imaging measures have shown consistency with clinical symptom progression and with Braak staging [5, 6], an accurate rendering of the 3D spatiotemporal profile of tau and A*β* at the micron scale has not been achieved [9, 10]. The principal challenge has been integrating the 2D sparse measurements of histology, which are direct measures of disease, with the MRI 3D markers, which are at much lower in-plane resolution. Consequently, these imaging measures have tended to emphasize MTL regions, including the entorhinal cortex (ERC) and hippocampus. These measurements, have not, however, been linked directly to micron level patterns of tau and A*β* pathology–the hallmark findings of AD.

The amygdala is one region that has undergone relatively limited study in AD [14] compared to its adjoining regions of the ERC and hippocampus in the medial temporal lobe (MTL) highlighted by Braak [5]. Neuropathological and connectivity-based studies suggest a role for the amygdala in neurodegenerative diseases such as Agrophylic Grain Disease (AGD) [15], Lewy Body dementia [16], and AD [17] with the inclusion of misfolded proteins and the amygdala’s strong connectivity patterns to areas of the hippocampus, basal ganglia, and basal forebrain [17]. Imaging studies from our group and others have observed cross-sectional differences in amygdala volume between cognitively normal individuals, those with MCI, mild AD dementia patients and later stage AD patients [18, 19, 13, 20]. These shape differences complement recent findings in tau PET illustrating cross-sectional differences in tau load in areas of the rostral MTL including ERC, TEC, and amygdala, in early stages of AD [21]. Two studies have also shown correlation between neuropsychiatric symptom severity and amygdalar atrophy in AD [22, 14]. These findings, together with emerging evidence of the neuropsychiatric syndromic complex known as Mild Behavioral Impairment (MBI) amongst individuals prior to the onset of dementia [23, 24] suggest amygalar changes early in the disease course may be related to the emotional and behavioral syndrome of MBI. Consequently, assessment of amygdala atrophy via modes as MRI holds promise as a biomarker for early diagnosis and management of AD. However, the exact timing and rate at which this atrophy occurs have not been well established as in other regions such as the ERC [25], but which is necessary for development of an appropriate biomarker.

To assess these rates, we compute both ERC thickness and amygdalar volume over time in MRI scans from a subset of the Alzheimer’s Disease Neuroimaging Intiative (ADNI) subjects in one of three categories: (1) stable controls (CON), (2) control subjects who progressed to MCI (PRE), and (3) MCI subjects who progressed to AD (MCI). Using an established approach of longitudinal diffeomorphometry [26], we estimate percent atrophy per year in each structure for each subject by fitting each subject’s series of scans to a smooth trajectory to extract signal from noise.

We subsequently link both ERC and amygdalar atrophy rates as MRI-based measures at a millimeter scale to patterns of AD pathology. We recently developed a method, referred to as Projective LDDMM, which facilitates the integration of multi-scale, multi-modal data into a single coordinate system [27]. Here, we use this approach to reconstruct 3D distributions of neuropathology within the MTL of advanced cases of AD, with a focus on comparing relative load of pathology in the ERC and amygdala compared with other regions of the hippocampus. As tau has exhibited stronger predisposition over A*β* for segregating to particular brain regions (ERC, CA1, subiculum) and layers (superficial) of cortex in AD [5], we use machine-learning based methods to detect neurofibrillary tangles (NFTs) from histological images. We quantify these detections in the space of histology images as measures of NFT density (per cross sectional tissue area) and carry these measures to the space of high field 3D MRI and the Mai Paxinos Atlas [28] using the transformations we estimate via Projective LDDMM. Using manual delineations on high field ex vivo MRI of subregions in the MTL and within both the amygdala and ERC, we quantify relative pathology within and among substructures for further corroboration of manifest atrophy rate with AD pathology.

## 2. Material and Methods

### 2.1. Subject Selection and Image Processing

#### 2.1.1. Longitudinal Data Selection

The subjects analyzed in this study were selected from the ADNI database (adni.loni.usc.edu) according to the criteria used in our previous studies [13, 25]. All subjects were required to have at least three 3T MR scans and at baseline, a diagnosis of stable neurocognition (NC), a CDR score [29] of 0, and normal range performance on the Logical Memory Subtest of the Wechsler Memory Scale [30] according to education adjusted norms. For segmentation and computation of cortical thickness in the entorhinal cortex (ERC), all subjects were also required to have an anteriorly continuous collateral sulcus, as detailed in [25]. Subgroups of stable NC, NC progressors to MCI, and MCI progressors to AD dementia, referred to as “CON”, “PRE”, and “MCI”, respectively, were delineated based on diagnosis at clinical follow-up. Table 1 summarizes the demographics of each of these three groups. Note that the accelerated scans of the ADNI 3 protocol were not included in this study.

**Table 1:** Demographics of three subgroups of ADNI dataset. Statistics reported as mean ± standard deviation where appropriate. Time of diagnosis reported as onset of MCI in PRE group and onset of dementia in MCI group.

| Category | CON | PRE | MCI |
| --- | --- | --- | --- |
| Sample Size | 33 | 16 | 18 |
| Baseline Age (years) | 72.3 ± 5.5 | 74.6 ± 5.2 | 72.8 ± 6.6 |
| Sex (% female) | 45.5 | 75.0 | 61.1 |
| # of scans | 3.9 ± 0.2 | 4.0 ± 0.6 | 3.8 ± 0.4 |
| Scan period (years) | 2.3 ± 0.7 | 1.9 ± 0.9 | 1.8 ± 0.4 |
| Time of Diagnosis<br>(years since baseline) | — | 4.1 ± 2.6 | 2.7 ± 1.8 |

**Table 2:** Donor demographics and pathological staging for ex vivo brain samples.

| Category | Subject 1 | Subject 2 |
| --- | --- | --- |
| Age | 93 | 87 |
| Gender | M | F |
| Clinical Diagnosis | Dementia | Dementia |
| Braak NFT stage | VI/VI | VI/VI |
| CERAD Neuritic Plaque Score [31] | B | B |
| TDP-43 | - | + |
| Pathologic Diagnosis [7] | High level AD pathologic change; multiple cerebral infarcts | High level AD pathologic change |

#### 2.1.2. Ex Vivo Specimen Preparation and Imaging

Brain tissue samples for ex vivo analysis were prepared from two cases of advanced AD by the Johns Hopkins Brain Resource Center. One hemisphere from each sample was reserved for neuropathological staging and diagnosis. The two samples examined in this study had clinical diagnoses of advanced AD and Braak stage VI pathology [6], indicative of high levels of tau pathology throughout the MTL. Detailed demographics and pathological staging of each sample analyzed are summarized in Table 2.1.2.

The other hemisphere was immersion fixed in 10% buffered formalin prior to dissection. A portion of the MTL extending from the temporal pole to the hippocampal tail was excised in 3 contiguous blocks of tissue, sized 20-30 mm in height and width, and 15 mm rostral-caudal. Each block was imaged with an 11T MR scanner at 0.125 mm isotropic resolution by the Mori lab at Johns Hopkins. Subsequently, the blocks were serially sliced into 10 micron thick sections, with sets of 5-6 sections taken approximately every 1 mm. Each block yielded between 7 and 15 sets of sections. The first section from each set was stained with PHF-1 for tau tangle detection. Remaining sections in each set were reserved for calibration of NFT detections between brain samples (see Section 2.3.2) or Nissl staining for confirming 3D MRI segmentations (see Section 2.1.3). All stained sections were digitized at 2 micron resolution.

#### 2.1.3. Regional Segmentations

Manual segmentations of MTL subregions were delineated on clinical (3T) and ex vivo (11T) MRI by a team of two individuals (CC and EX) guided by a neuroanatomist (MW). All segmentations were drawn with Seg3D version 1.13.0 [32]. Prior to segmentation, brightness and contrast on the MRI was adjusted for better visualization. A brush size of 1 voxel was selected for precision. Segmentation was primarily done on the coronal plane, while the axial and sagittal plane were used to clarify borders.

In clinical 3T scans, ERC and TEC were delineated following the protocol used previously [13, 25, 26]. Visible anatomical landmarks were defined to match described cytoarchitectonic borders [33, 34]. Anteriorly, the ERC and TEC extended 4mm beyond the hippocampal head. Posteriorly, the ERC and TEC extended 2 mm beyond the gyrus intralimbicus (GI) [33]. Medially, the ERC extended to the edge of the visible gray/white matter boundary. Laterally, the TEC extended to the deepest portion of the collateral sulcus. Finally, the border between lateral ERC and medial TEC was defined at the medial extent of the collateral sulcus [33].

The amygdala was segmented coronally slice by slice from posterior to anterior following a novel protocol based on intensity contrast and comparison with anatomical landmarks identified for the hippocampus [35]. Posteriorly, the region of dark contrast superior to the hippocampus, representing the inferior horn of the lateral ventricle, was used to delineate the anterior hippocampus from the posterior start of the amygdala. At that level, the first set of voxels marked as amygdala were those exhibiting an intermediate gray level of intensity, rather than white matter contrast as appears posteriorly. Over its antero-posterior extent, the amygdala was defined as an area with even intensity contrast that increased in size anteriorly. The anterior border was demarcated by the disappearance of contrast, which occurred, on average, 11.8 mm ± 2.0 mm from the marked posterior border. For consistency, one row of empty voxels was maintained between the amygdala and ERC and TEC.

In ex vivo 11T scans, MTL subregions included amygdala, ERC, CA fields, subiculum, presubiculum, parasubiculum, and dentate gyrus. Segmentations were drawn following manual alignment of individual block MRIs into a single MRI using an in-house alignment tool. Boundaries for each region were defined based on intensity differences and published MR segmentations [36, 37, 38], following a novel protocol [35]. Where available, Nissl-stained histology sections were overlaid with MRI for validation of the intensity-based protocol. For assessment of NFT density distribution within the amygdala, the amygdala segmentation in a control brain sample was subdivided into five individual regions based on visible intensity differences in 11T images, combined with anatomical markers and previously published delineations [39, 40]. Regions include basolateral (BLA), basomedial (BMA), cortical, medial, and central (CMA), lateral (LA) and periamygdaloid (PA) nuclei.

For evaluating accuracy of alignment between digital pathology and MRI, as done previously [27], all digital pathology images from a single brain sample were segmented into corresponding MTL subregions. These segmentations were defined on 4x-downsampled histology images at a resolution of 32 microns. The protocol for segmentation of 2D images used previously published delineations of MTL subregions based on cytoarchitectonic characteristics [39, 40, 41, 42, 43, 44].

### 2.2. Morphometric Measuring of Atrophy Rate

#### 2.2.1. Smoothing via Longitudinal 3D Diffeomorphometry

Following the approach developed by Tward and Miller [26], we estimate the flow of a template image, *I*_temp_, through the series of each patient’s images, 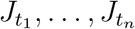 observed at times *t*_1_, …, *t*_*n*_ to compute atrophy over time for each subject. This yields a smooth flow over time that effectively acts as a filter over the image series, eliminating variance in volumetric calculations due to imaging parameters such as head placement, specific machine, etc. We define the estimation of the template’s flow through the time series:

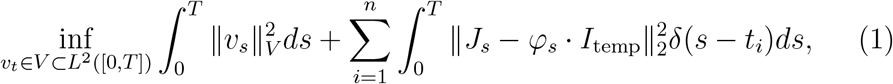

for 0 ≤ *t*_1_ ≤ *t*_*n*_ ≤ *T*.

In our problem, *I*_temp_ is not fixed but is estimated for each patient via a flow of a population template to the patient’s geodesic trajectory at a certain time, *t*^∗^, which is estimated in tandem with the trajectory, with *t*_1_ ≤ *t*^∗^ ≤ *t*_*n*_ [26]. Consequently, the population template is mapped to each target subject in our ADNI cohort and then forwards and backwards in time according to a smooth geodesic trajectory. This formulation ensures a regularization cost over the whole geodesic trajectory for each target to achieve similar smoothing over the length of the trajectory. Finally, we generate image series for each subject smoothed over time by sampling the estimated flow of our template through each subject’s time series at the corresponding times of the subject’s original scans. This process is summarized in Figure 1, where amygdala surfaces for one subject at each of four time points are shown in blue, overlaid with corresponding surfaces of the template’s flow sampled at these same time points. Estimated volume measures compared with original measures exhibit smoothing as a consequence of this method.

**Figure 1:**
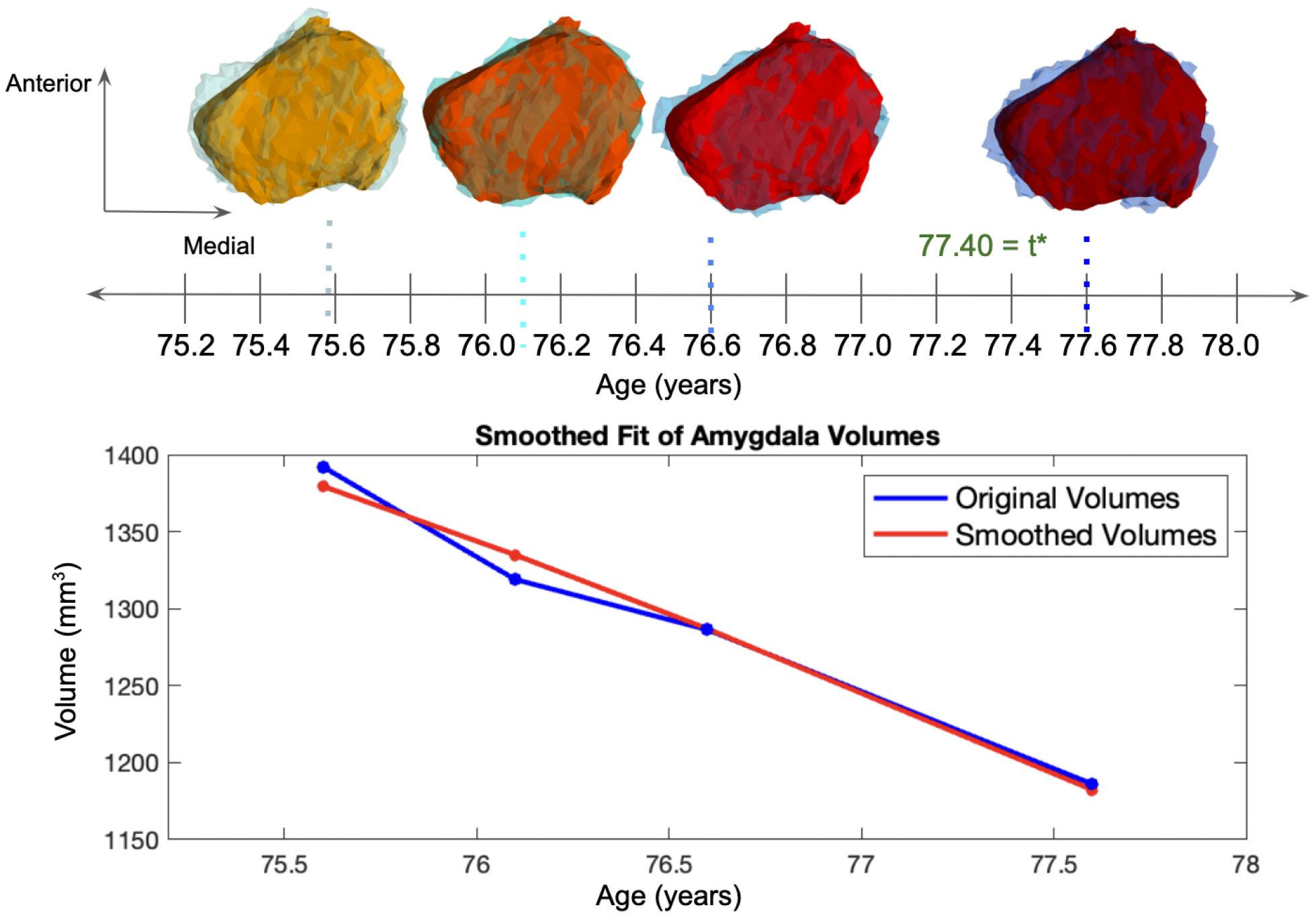
Smoothed flow of amygdalar surface for one subject in preclinical converter cohort. Estimated surfaces sampled along the template’s flow at each of four time points shown in red overlaying original surfaces in blue at the top. Measured volumes of original surfaces and estimated smoothed surfaces plotted in bottom figure.

Here, we use two different population templates for mapping to combined ERC and TEC versus amygdala. As in previous work, we estimate a template ERC plus TEC surface mesh from the set of all our ERC and TEC surface meshes from each subject of each group at each time point [45]. To examine atrophy in the amygdala, we generate a template surface mesh from a high field ex vivo MRI (0.125 mm resolution) of a 27 year old male without AD pathology. This sample was prepared according to the protocol described in Section 2.1.2, and both regional and subregional delineations of the amygdala were drawn on this MRI following the protocol described in Section 2.1.3.

#### 2.2.2. Amygdala Volume Measurements

Smoothed estimates of amygdala volume (and ERC thickness) following longitudinal shooting (see Section 2.2.1) were computed from 3D surfaces of the template resampled at each target time point in a subject’s time series. 3D surface mesh renderings of the amygdala and the combined ERC and TEC were generated for each subject at each time point using restricted Delaunay triangulation [46]. We define a triangle family as a collection ***x*** = (*x*_*i*_, *i* ∈*I*) of distinct points in *R*^3^ together with a family of 3-tuples, *C* = (*c* = (*c*_1_, *c*_2_, *c*_3_) ∈ *I*^3^) of indexes such that the cells

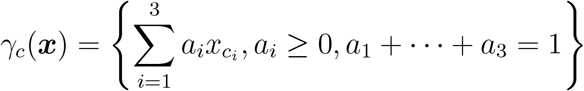

have non-empty interior with positive orientation, i.e., their area is

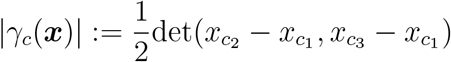

where the term in the right-hand side is positive. The cell centers *m*_*c*_ are

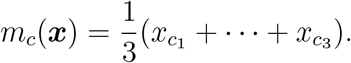

Amygdalar volumes were computed from triangulated surfaces by summing the signed individual volumes of tetrahedrons formed by each cell together with the origin:

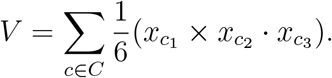

Atrophy rate per subject was measured in the amygdala as change in volume per time (mm^3^/year). Rates of change were estimated per subject for a subject’s entire time series via least squares fitting for each of the three groups. For comparison between subjects, rates were computed as a percent change in volume from the starting measure in each series of time points.

#### 2.2.3. ERC plus TEC Thickness Measurements

Vertex-wise thickness of combined ERC and TEC surfaces was computed following the established approach of Ratnanather et. al [47], as used in our previous work [13, 25]. Surfaces were cut into an outer (pial) surface and inner (gray/white matter boundary) surface using an in-house manual tool. Diffeomorphic transformations of the inner surface to the outer surface were estimated with LDDMM with the added constraint that the deforming inner surface flowed normal to itself at each sampled point in the flow. Vertex-wise thickness was estimated as the distance each inner surface vertex traveled over the course of the diffeomorphic flow to the outer surface. Composite ERC and TEC thickness measures for each subject at each time point were estimated as the average vertex-wise thickness from the set of vertices in the inner surface correspondingly labeled as ERC or TEC respectively.

Rates of change in the ERC and TEC were computed as change in thickness per time (mm/year). For each subject, average atrophy rate was estimated by least squares fitting to the subject’s entire time series of smoothed measurements. As with amygdalar atrophy rates, ERC plus TEC rates were computed as a percentage change in thickness per year from baseline measurement.

### 2.3. Ex Vivo Tau Reconstructions

#### 2.3.1. 3D Reconstruction of 2D Digital Pathology

Sets of 2D digital histology images at 2 *µ*m resolution were mapped to the space of 3D high field ex vivo MRI at 0.125 mm resolution following our approach, Projective LDDMM, as detailed in [27]. Following the random orbit model of computational anatomy [48], each histology section is modeled as a Gaussian random field with mean field modeled as an image in the orbit of a template *I*_temp_ under the group of diffeomorphisms, *I* ∈ ℐ = {*I* = *φ* · *I*_temp_, *φ* ∈ *G*_diff_}. To accommodate differences in geometric dimension of digital histology images (2D) and MRI (3D), the mean of each field is given as the result of applying a projection operator to generate the targets *P*_*n*_ : *I* → *J*_*n*_ = *P*_*n*_*I* + noise, *n* = 1, …, *N*, with observed target images, *J*_*n*_, modeled as conditional Gaussian random fields with mean fields, *P*_*n*_*I*. The projections *P*_*n*_*I*(*y*) are intrinsic to the histology process prescribing the weight an image value *I*(*x*) at location *x* contributes to the target image value *J*(*y*) at *y*. For serial sectioning at 1 mm intervals along the anterior-posterior axis of the brain:

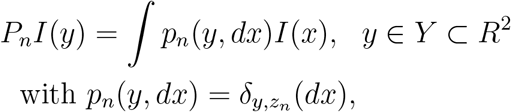

where the Dirac delta measure, 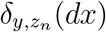, assigns nonzero measure only to the image value *I*(*x*) for *x* = (*y*^(1)^, *y*^(2)^, *z*_*n*_), effectively restricting the template image to that within the image plane of target *J*_*n*_ at location *z*_*n*_ along the anterior-posterior axis of the brain. The tissue sectioning in histology introduces additional deformation modeled as 2D transformations *ϕ*_*n*_ in the 2D image plane, independent from section to section *n* = 1, …, *N* :

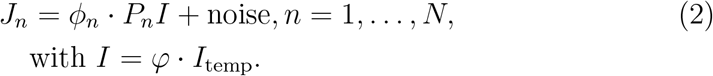

Here, each *ϕ*_*n*_, is modeled as a diffeomorphism. We estimate *φ* and each *ϕ*_*n*_ as the flow of time-varying velocity fields, following the approach of Large Deformation Diffeomorphic Metric Mapping (LDDMM) of Beg et. al [49]. We alternately optimize our cost function (3) for *φ* and each of the *ϕ*_*n*_ while holding the other fixed, as described in [27]. Estimation of *φ* is explicitly given for optimizing over *v*_*t*_, *t* ∈ [0, 1] to:

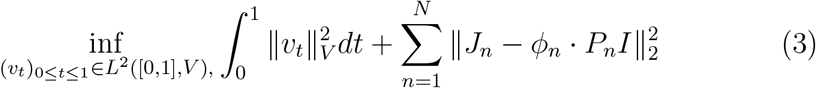

with ∥ · ∥_*V*_ a norm defined over a space *V* of smooth time varying velocity fields and ∥ · ∥_2_ the integral square norm. Estimation of each *ϕ*_*n*_ proceeds via estimation of velocity field (*u*_*t*_)_0≤*t*≤1_ ∈ *L*^2^([0, 1], *V*) with 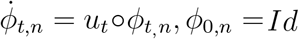.

To resolve differences in the multi-scale resolution and contrast of the digital histology micro-scale images in relation to the tissue scale MRI, we expand the histology images using a Scattering Transform [50]. The Scattering Transform is comprised of an alternating sequence of convolutions with wavelets and non-linear modulus operations (i.e. modulus) that generate a sequence of multi-scale images 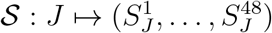, which reflect the tissue characteristics originally encoded at the high resolutions of digital pathology. We use PCA to build a 7-dimensional linear predictor from the scattering images with a constant image [27], which approximates an MR contrast image of histology as a function of the predictor *J*_*n*_(***α***), ***α*** = *α*_1_, …, *α*_7_. The optimization problem simultaneously solves for the diffeomorphism transforming the template onto the target histology with (3) and the low-dimensional linear predictor, with estimation of ***α***_***n***_ following least squares minimization: 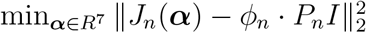 [27].

For distortion extending to tears, tissue folding, and other artifacts, we introduce a set of latent variables in the context of Gaussian mixture models, assigning each pixel location in the image plane to one of three classes: foreground tissue, background tissue, or artifact, as described in [51]. Estimation of latent variable values together with each *ϕ*_*n*_ is achieved with the Expectation-Maximization (EM) algorithm [52], resulting in a cost function with matching term weighting each location in the image plane according to its iteratively estimated posterior probability of being foreground tissue [27]. The high field MRI was collected in three blocks.

To integrate measures across blocks, we used an in-house tool to rigidly align each set of three MRI blocks. The Mai Paxinos Atlas [28], was rigidly aligned to a surface rendering of the complete hippocampal geometry of each brain sample manually to a surface rendering of the Mai hippocampus. The findings reported in Section 3 reflect the coordinate system of the Mai atlas.

#### 2.3.2. Neurofibrillary Tau Tangle Density Measures

We quantified tau pathology from digital histology images as number of neurofibrillary tau tangles (NFTs) per cross-sectional tissue area. Measures of NFT density were computed over each section following the approach described previously [27]. For each brain sample, a UNET [53] with architecture specified in Appendix A, was trained on a subset of patches of background and foreground tissue with NFTs manually annotated. Resulting probability maps predicted the likelihood of each pixel’s being part of an NFT. Probability maps were then segmented using an available implementation of the Watershed algorithm [54] to delineate individual NFTs. Each histology image was segmented into foreground tissue and background using Otsu’s method [55], and final measures of NFT density per slice, region, or subregion were computed by taking a sum of NFTs within foreground tissue divided by the square mm area of foreground tissue in the region of interest.

#### 2.3.3. 3D Reconstruction of NFT Densities

Distributions of tau pathology in 3D were reconstructed per brain sample following estimation of geometric mappings between histological images and 3D MRI (see Section 2.3.1). Estimated NFT densities within each image plane were modeled as features, *f*_*i*_, associated to “particles” at locations, *x*_*i*_, in a regular voxel grid, using a measure-based framework, as described in [56]. Weights, *w*_*i*_, reflecting the square mm area of tissue captured by the *i*th particle at each position *x*_*i*_ were associated with the particle measures (mathematical measures, see [56]) and carried to the Mai coordinate space, together with MRI and associated MTL segmentations, following the prescribed action of diffeomorphisms on measures [56].

In the Mai coordinate space, measures were pooled across histological sections and resampled within the dense volume. Physical and feature space were transformed independently, as enabled by the decomposition of measures into a physical density (cross-sectional tissue area (mm^2^) per unit of 3D space) and conditional feature probability (number of 2D detected NFTs in unit of 3D space given its physical density). Spatial resampling to a regular grid within the volume of the Mai atlas at a resolution of MRI was achieved with a Gaussian kernel, assigning a fraction of each particle measure’s weight, *w*_*i*_, to each new particle *x*′ with new weight, 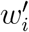 in the resampled space. Feature values for each new particle *x*′ were computed as the expected first moments according to the empirical distribution defined by the spatial reassignment of weights.

#### 2.3.4. Laplace-Beltrami Resampling to Surfaces

In addition to resampling within the dense volume, resampling along the surface manifold of MTL subregions was achieved with a nearest neighbor kernel, “projecting” NFT density measures to the boundary of each surface by assigning the entirety of each particle’s tissue area within a given region at high resolution to a single particle within the manifold. Smoothed NFT densities were estimated as the ratio of smoothed NFT counts 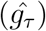 to smoothed cross-sectional tissue areas 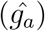 following expansion of each vertex defined function *g*_*τ*_ (*x*_*i*_) := *f*_*i*_, *g*_*a*_(*x*_*i*_) := *w*_*i*_ in the Laplace-Beltrami basis, as described previously [57, 27]. Specifically, for each of *g* = *g*_*a*_(·) and *g* = *g*_*τ*_ (·), we compute:

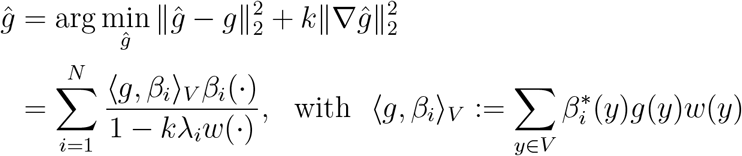

where ℬ := {*β*_1_, …, *β*_*N*_} is a basis for the Laplace-Beltrami operator, *w*(·) defines a weight for each vertex based on partitioning the surface area of adjoining faces, and *k* is the smoothing constant, with value of 2 in the results presented in Section 3.

## 3. Results

### 3.1. Amygdalar and Entorhinal Atrophy Rates

Following estimation of a smoothed trajectory over time for each subject via longitudinal diffeomorphometry (see Section 2.2.1), this trajectory was resampled at each subject’s original scan time points. The results of this resampling are plotted for each subject in Figure 2. Average atrophy rates for ERC thickness and amygdala volume were computed from the smoothed measures for each series of time points for each subject via least squares fitting. Scan periods ranged from 1 year to 5.1 years for sets of time points, and each subject had between 3 and 7 scans. In both entorhinal and amygdalar regions, MCI converters and preclinical converters exhibited higher atrophy rates than stable controls. Mean atrophy rates of the ERC plus TEC were 1.1, 3.8, and 7.0% thickness loss per year, and of the amygdala, were 1.5, 6.8, and 11.6% volume loss per year for stable controls, preclinical converters, and MCI converters, respectively.

**Figure 2:**
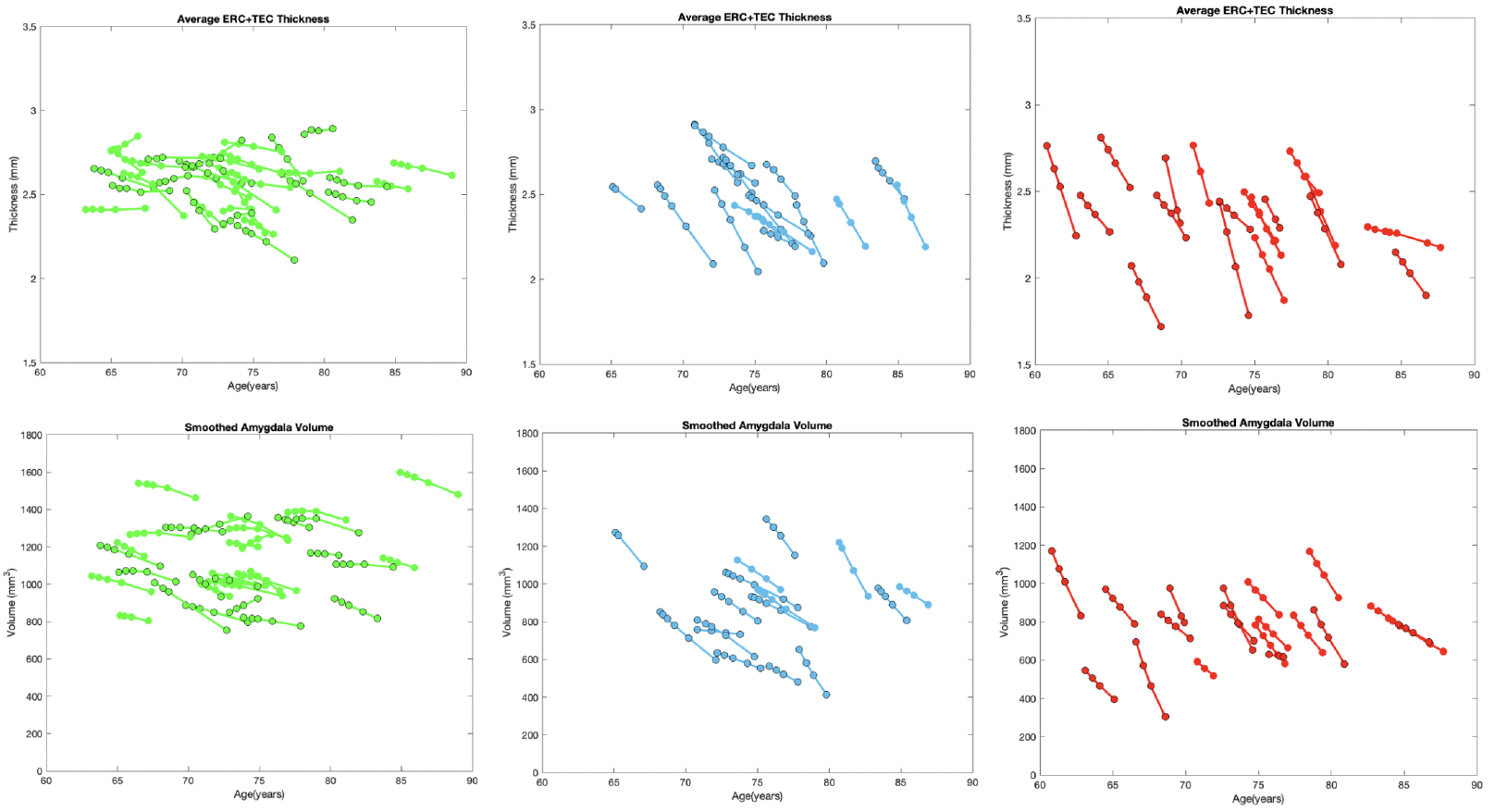
Smoothed measures of ERC plus TEC thickness (top) and amygdala volume (bottom) plotted against subjects’ ages at each scan time point. Stable controls in green (left), preclinical converters to MCI in blue (middle), and converters to AD in red (right). Female patients outlined with black circles. Mean atrophy rates (± standard deviation) of ERC plus TEC are 1.1 ± 1.6%, 3.8 ± 1.9%, and 7.0 ± 3.4% thickness loss per year in controls, converters to MCI and converters to AD, respectively. Mean atrophy rates of amygdala are 1.5 ± 2.0%, 6.8 ± 4.1%, and 11.6 ± 5.8% volume loss per year.

To assess the potential of amygdalar atrophy rate, measured via 3T MRI, as a biomarker for AD, we compared distributions of estimated atrophy rates in controls to those of preclinical and MCI converters via ROC analysis. Distributions are illustrated as histograms with estimated kernel density estimates for each clinical group (Figure 3). ROC analysis yielded an optimal threshold of discrimination between controls and both preclinical and MCI converters at −4.7% volume loss per year, with sensitivity of 0.91, specificity of 0.91, and AUC of 0.96.

**Figure 3:**
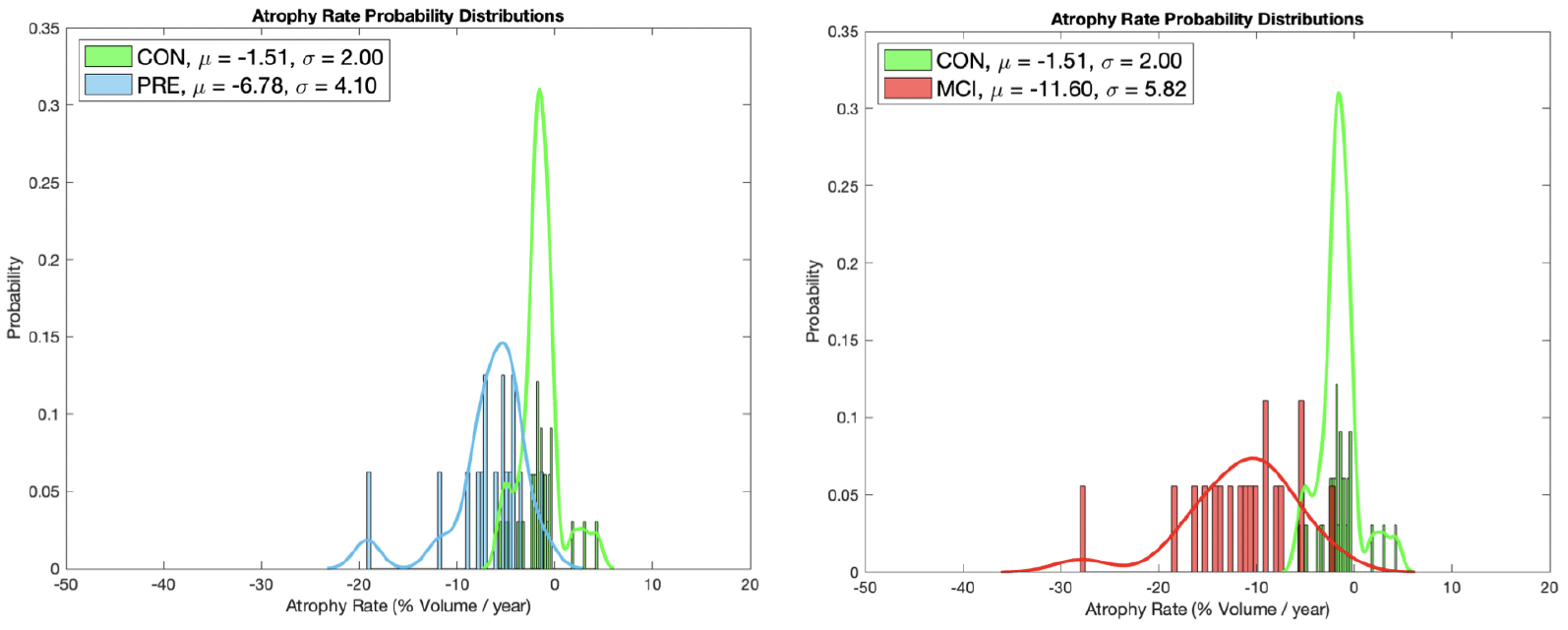
Distribution of amygdala atrophy rates estimated from each set of scan time points per subject. Distributions of controls (green) shown overlaid with distributions of converters to MCI (left) and converters to AD (right). Mean and standard deviation are 1.5 ± 2.0%, 6.8 ± 4.1%, and 11.6 ± 5.8% volume loss per year for controls, converters to MCI, and converters to AD, respectively.

### 3.2. Geometric Reconstructions of Postmortem Brain Samples

Two sets of 35 digitized histological sections stained with PHF-1 were aligned to 3D high field MRI blocks of corresponding MTL tissue following the approach described in Section 2.3.1. Alignment accuracy was evaluated by comparing 2D segmentations of MTL subregions on histological images to 3D segmentations of corresponding regions deformed to 2D. Dice Score and 95th Percentile Hausdorff distance for each region on each slice of one brain were computed with average overlap scores of 0.85, 0.82 and average 95th percentile Hausdorff distance of 1.886 mm and 1.039 mm for the amygdala and ERC, respectively. Positions of each section within each block are shown for one brain sample in Figure 4 with manual segmentations of MTL subregions illustrated across the rigidly aligned MRI blocks. Following alignment of 2D histological sections to 3D MRI, both sets of images were transported to the space of the Mai Paxinos atlas [28] with estimation of rigid alignment between hippocampal surfaces for each brain sample and that of the atlas. Figure 5 exhibits surface renderings of select MTL subregions for one brain sample in the coordinate system of the Mai Paxinos atlas. Snapshots of the atlas show coronal slices in the area of the anterior MTL intersecting with select histological slices.

**Figure 4:**
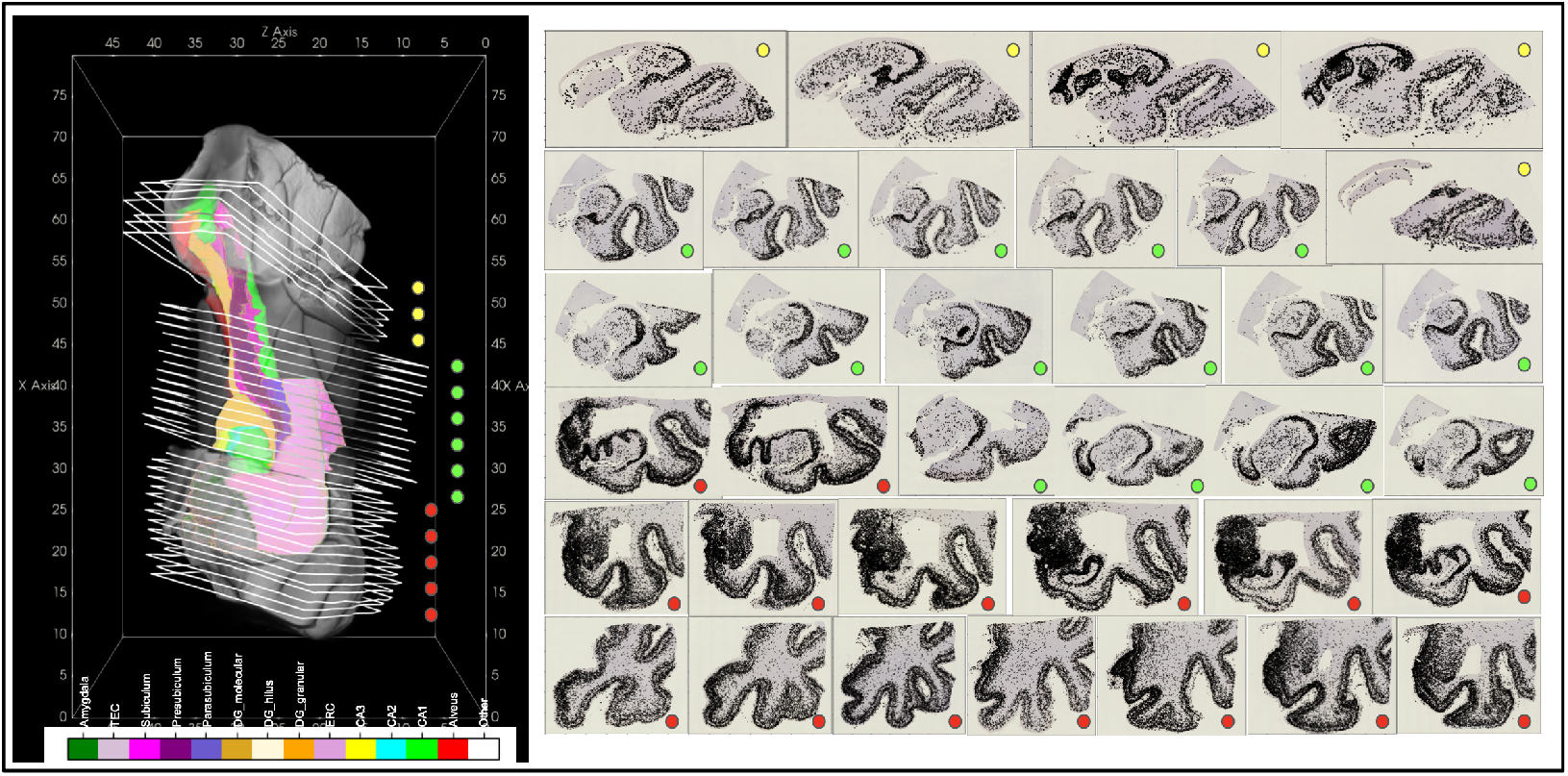
Complete sets of digitized PHF-1 stained histology sections for 3 blocks of MRI for an AD brain sample. 3D MRI shown with manual segmentations of MTL subregions (left). Boundary of each histological section on right shown in white in position following transformation to 3D space (left). Detected NFTs plotted as black dots over each histology slice (right).

**Figure 5:**
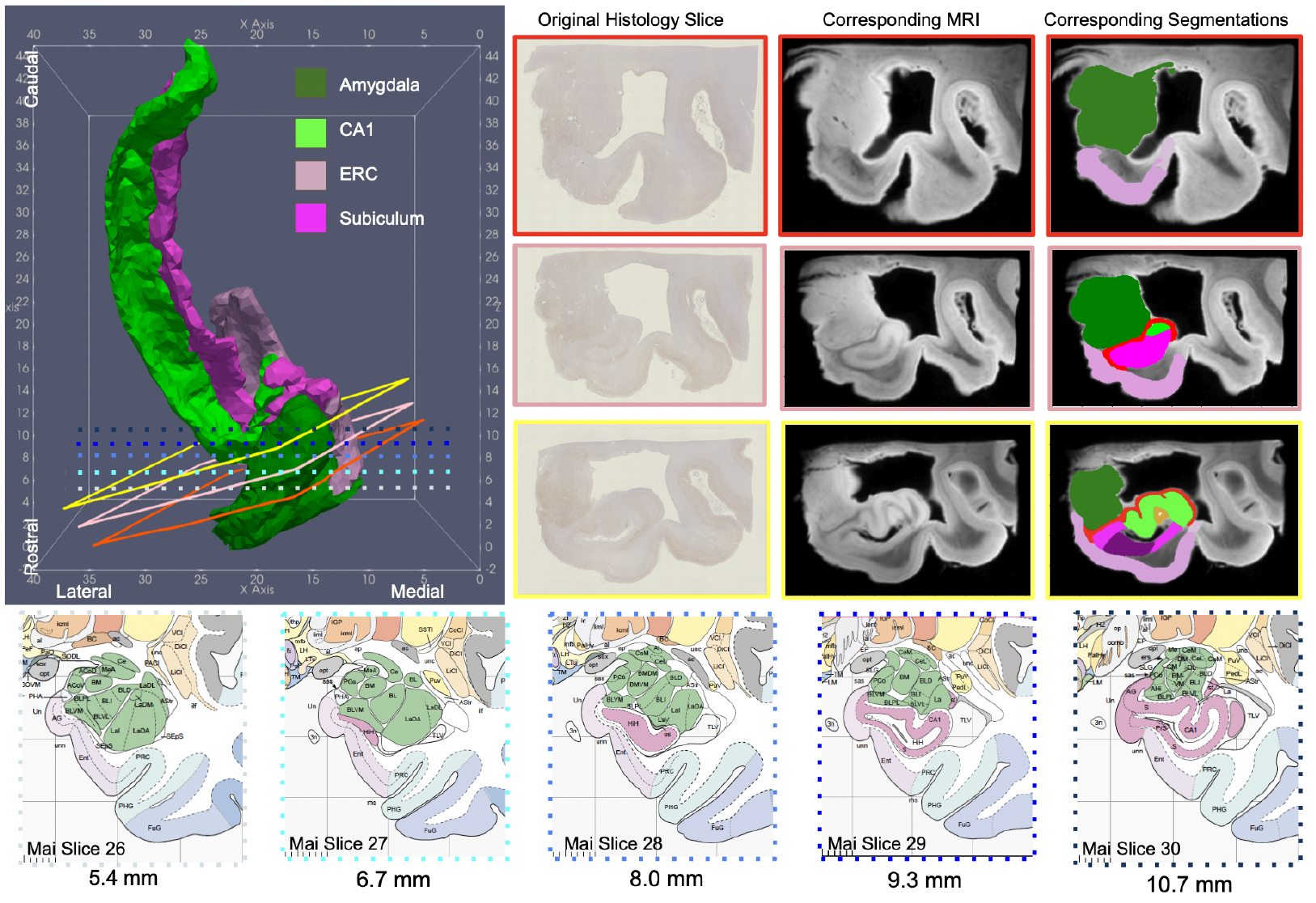
3D Reconstruction (left) of 4 MTL subregions for an AD brain sample in the coordinate space of the Mai Paxinos Atlas. Example section of histology and corresponding MRI slice (right) shown at approximate location of intersecting coronal planes taken from the pages of the Mai Atlas (bottom).

### 3.3. NFT Densities in Advanced AD

NFTs were detected on each histological image for each brain sample with separate UNETs trained independently on a subset of training data for each sample. We evaluated the accuracy of UNET-predicted per-pixel probability of tau using 10-fold cross validation. For the two brain samples, mean AUC was estimated at 0.9860 and 0.9873, and mean accuracy at 0.9729 and 0.9546, respectively (see Appendix A for details). We evaluated the accuracy of discrete NFT detections following segmentation with the watershed algorithm by direct comparison to a set of manually annotated image patches reserved for validation. Total counts of NFTs annotated per patch varied consistently from patch to patch with total counts of NFTs detected. Exact measures provided in Appendix A.

2D measures of NFT density were carried to the space of the Mai Paxinos atlas, together with MRI and manual segmentations of the MTL. NFT density measures were pooled across sections and resampled at 0.2 mm resolution within the coordinate space of the Mai atlas (see Figure 6). Deformation of 3D MTL segmentations to 2D histology images enabled assignment of NFTs into MTL subregion. NFT densities per MTL subregion were computed overall (see Figure 7) and over the surface boundary of each region (see Figure 6), following smoothing with the Laplace-Beltrami operator, as described in Section 2.3.4.

**Figure 6:**
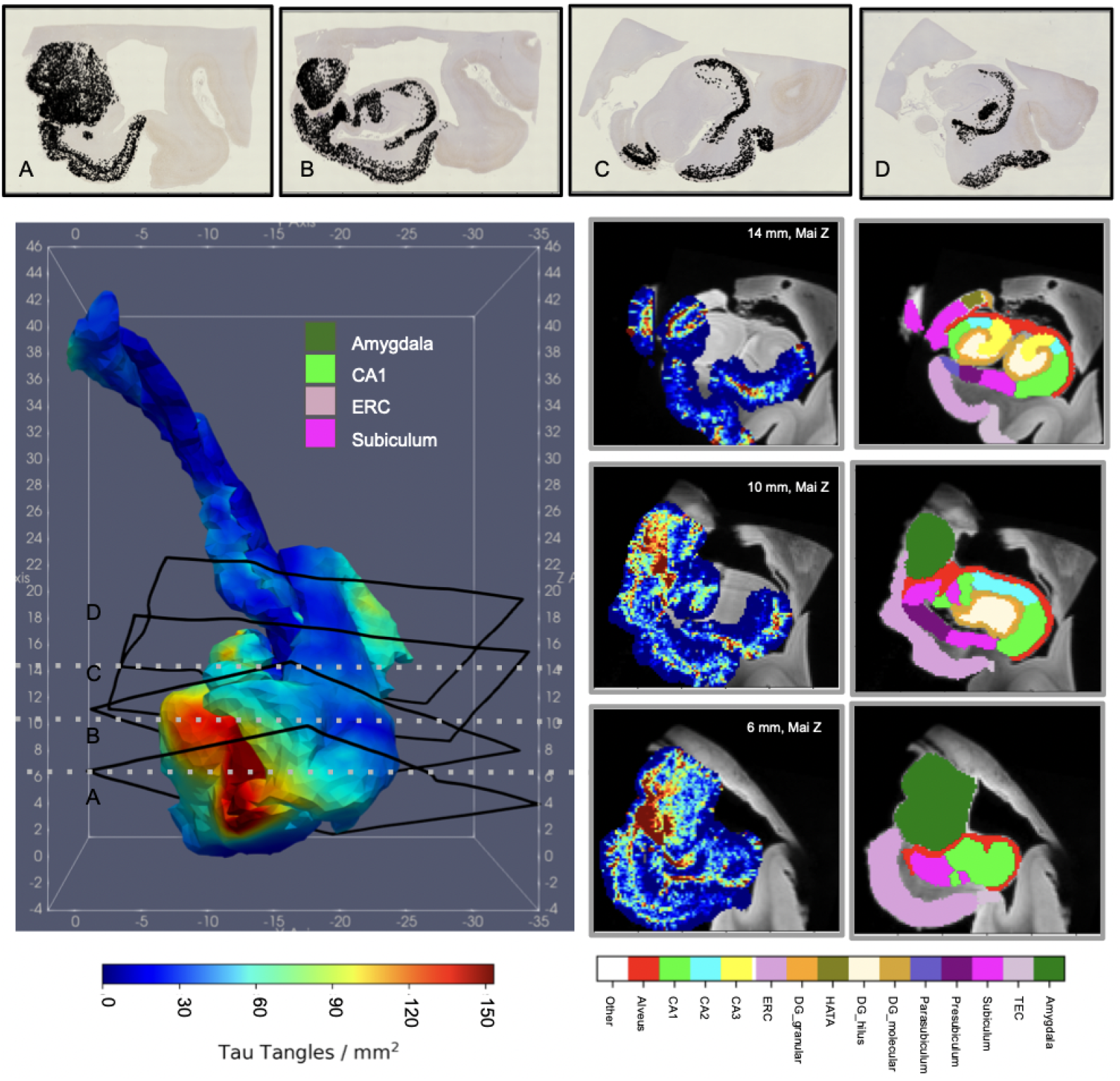
Distributions of NFT densities computed from digital histology at 2 *µ*m (top) reconstructed in 3D for an advanced case of AD. Densities within a subset of regions of MTL (amygdala, ERC, CA1, and subiculum) sampled within the dense metric of the 3D MRI at 0.2 mm resolution (right) or projected to the surface of each structure and smoothed with the Laplace-Beltrami operator (left). Sections of MRI shown at 6, 10, and 14 mm, corresponding to coronal slices in the Mai Paxinos atlas.

**Figure 7:**
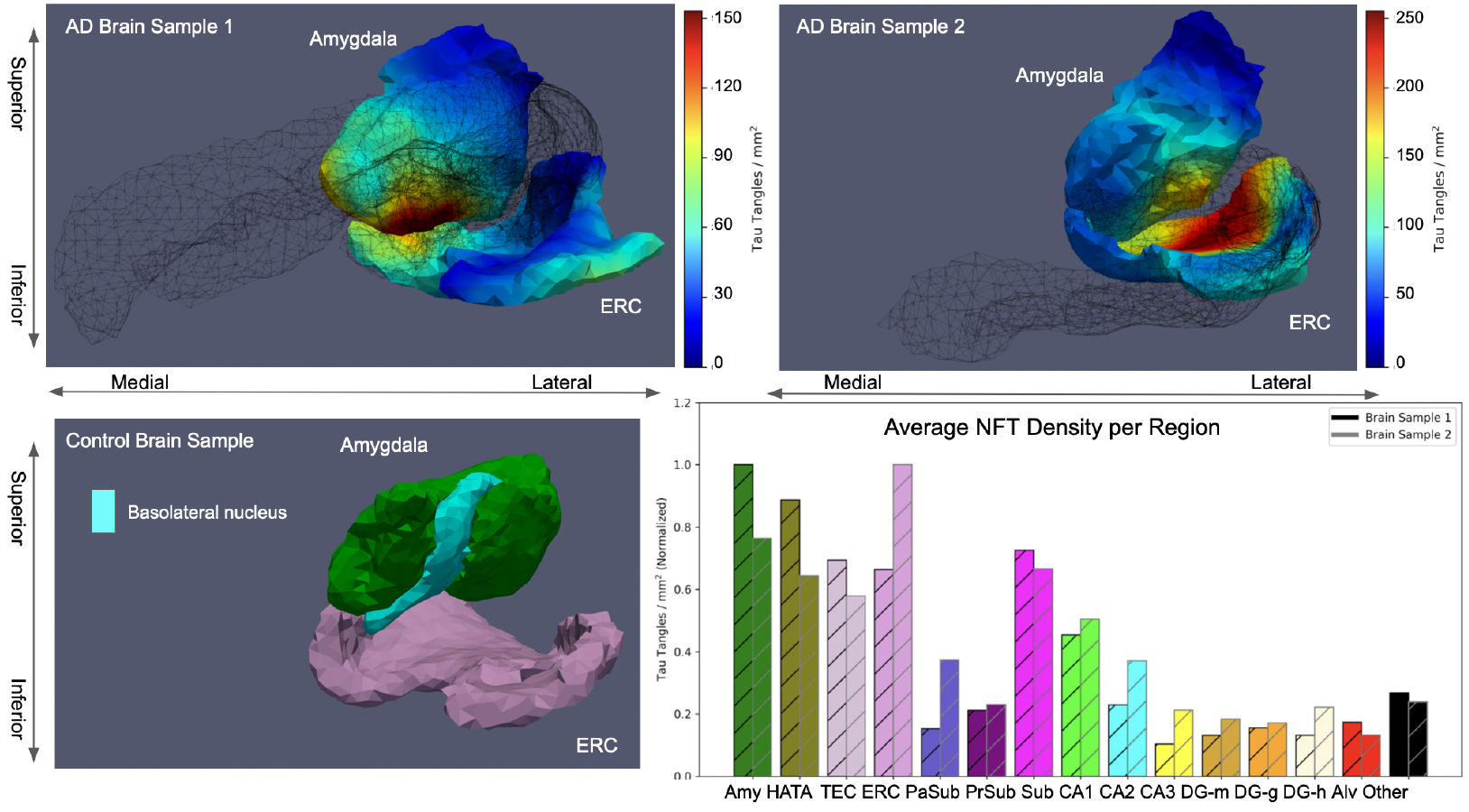
Posterior view of amygdala-ERC boundary in control (bottom left) and two AD brain samples (top). NFT densities for AD brain samples shown following projection to each surface and smoothing with the Laplace-Beltrami operator. Outline of CA1 and subiculum surfaces shown in black mesh. Basolateral amygdalar subregion delineation shown for control brain sample. Average NFT density within each MTL subregion normalized to maximum average for each AD brain sample (bottom right).

NFT densities are reported on different scales for each brain sample. To compare NFT densities across brain samples, subsets of 4-5 additional sections from each brain were selected in the rostral hippocampus at approximate locations of original sections. These two subsets were stained simultaneously to achieve consistency between them and the respective UNET trained on the original training data from each brain was used to detect NFTs in these sets of replicate slices. Ratios of NFTs detected on the original and new version of each section were computed, yielding average ratios of 2.7 and 27.2 for brain samples from subjects 1 and 2, respectively. These differences in level of detection speak to the effect that variation in staining intensity, timing, and handling of tissue samples has on absolute counts of NFTs. Therefore, to compare relative distributions of tau tangles between the two brain samples (see Figure 7), we normalized NFT density measures accordingly in each brain to the range 0 to 1. Average NFT densities per MTL subregion showed highest amounts of NFTs in amygdala, ERC, CA1, and subiculum for both advanced AD samples (see Figure 6,7).

Tau pathology localized not just *to* particular regions (e.g. amygdala, ERC, CA1, subiculum), but also within them. As illustrated in Figure 7, high densities of NFTs in amygdala and ERC concentrated particularly at the border between the two structures, with tau migrating to the inferior, medial boundary of the amygdala, particularly in its basomedial and basolateral regions. Highest NFT densities within the ERC, amygdala, and hippocampal-amygdala transition area (HATA) constitute a propensity of NFTs within the rostral third of the MTL.

## 4. Discussion

There are two primary findings from this research. First, these results demonstrate that the amygdala atrophies at a significantly faster rate in preclinical subjects compared with controls. In both the ERC and amygdala, our examined controls exhibited some level of atrophy (1.1% thickness loss and 1.5% volume loss, respectively). Likely an aspect of normal aging independent of disease, these atrophy rates nevertheless suggest that these control subjects, if they live long enough, might develop cognitive impairment given the accumulated atrophy of the ERC and amygdala. However, the differences between preclinical converters and controls in volumetric atrophy rate as measured by MRI from a series of 3-6 scans are comparable to those in thickness atrophy rate of the ERC plus TEC, with preclinical converters atrophying at a rate over 3x that of controls and MCI converters at a rate over 6x that of controls. In terms of onset, we have previously shown that the MTL as a grouped structure including ERC, TEC, and amygdala exhibits a changepoint nearly a decade before clinical time [12]. The findings here explicitly analyzing amygdala and ERC separately in preclinical and MCI groups indicate an early role for this MTL aggregate marker.

Second, these results demonstrate concordance between patterns of microscopic NFT pathology, specific to AD, and these macroscopic measures of atrophy through reconstruction, for the first time, of microscopic measures in the dense metric of MRI. Highest densities of NFTs not only in the ERC, but also in the amygdala, putatively suggest that the NFT pathology is underlying observed atrophy. Also, as MTLs from advanced cases of AD, these reconstructed distributions exhibiting highest end stage densities in ERC, HATA, and amygdala (see Figure 7) suggest that one may consider AD as a rostrally-oriented disease of the MTL.

The emergent role of the amygdala, here, is supported by our own work in shape diffeomorphometry [18, 19] as well as recent studies in MRI [14], tau PET [21], and tau pathology [58] that have seen similarly high levels of atrophy or tau pathology in the amygdala, particularly in early AD. Further work is needed to sufficiently delineate which areas of the amygdala might be more involved in early AD, but in both brain samples analyzed here, basolateral and basomedial regions of the amygdala, often classified together with the lateral nucleus as “core” amygdala [59], demonstrated higher tau densities as those adjacent to the ERC (Figure 7). Evidence of differential involvement of amygdalar nuclei has been cited in other neuropsychiatric diseases, such as Parkinson’s Disease [60] and depression [59]. Following [39], we are currently segmenting subregions of the amygdala in high field MRIs of MTLs from subjects with early and advanced AD to further assess specificity of tau pathology for particular amygdalar nuclei in the context of AD.

This work has both strengths and limitations. First, it is one of few studies integrating microscopic *and* macroscopic scales of analysis through digital pathology and MR imaging measures. Having developed the technology and mathematical infrastructure to integrate these two data types, we are poised to analyze other cohorts to corroborate the putative link between amygdala and ERC atrophy observed in MR imaging to underlying NFT pathology. Second, many previous clinical imaging studies from our own group as well as others [18, 19, 14, 21] have compared control and disease groups only cross-sectionally at a single time point. The longitudinal analysis here tracks subject specific measures over time. As such, it reduces variance inherent to the use of single measures to indicate disease status. Though the ERC’s early involvement in AD was established by Braak [5], the level and timing of the amygdala’s involvement remains an open question. Further longitudinal analysis will provide greater insight into the exact timing and progression of both ERC and particularly amygdala involvement in AD, as necessary for the development of biomarkers. Third, compared with other recent efforts to reconstruct distributions of tau pathology in 3D [58], we provide a quantitative metric of pathology (NFT density) which retains much of the precision found at the microscopic scales at which the pathology was identified. Lastly, manual segmentations of regions of the MTL in both MR images and ex vivo MRI offer an improvement over past efforts analyzing amygdalar atrophy and pathology in which segmentations were estimated from alternative templates [13, 58, 61].

Limitations of this study include both lack of data points and restricted analysis on data points available. Here, we present 3D reconstructions of MTL NFT pathology from only two brain samples, both with advanced AD. To build further support for the specificity of clinical imaging biomarkers such as MRI-measured ERC and amygdala atrophy rate in signature patterns of pathology in AD, we are currently reconstructing the distributions of both NFT and A*β* pathology in early, intermediate, and advanced stages of AD.

Additionally, atrophy rates were assessed for ERC and amygdala only in the left cerebral hemisphere whilst NFT reconstructions were computed for each of one left and one right hemisphere. Previous studies have suggested differences of involvement in right versus left amygdalas [62], and future work will be needed to confirm or deny any differences in pathology versus atrophy in each hemisphere. Finally, while this work aims to link pathology at the micron scale to MR imaging markers at the mm scale, emerging technologies in PET imaging of both tau and amyloid [63, 4] offer an opportunity for integration at an intermediate scale for both corroborating the results here and enabling the development of biomarkers across multiple modalities.

In conclusion, we have demonstrated the existence of significant atrophy in *both* the ERC and amygdala early in the course of AD. Measurable from a series of MR scans and putatively linked to high levels of AD specific pathology, atrophy rates in both ERC and amygdala present as viable biomarkers that may someday be used for both early diagnosis and management of AD.

## Appendix A. UNET Details

Individual NFTs were detected from digital histology images of tissue samples stained with PHF-1. Detection followed a two-step algorithm: 1) prediction of each pixel’s probability of being part of an NFT and 2) segmentation of probability maps into discrete connected components corresponding to NFTs. The architecture of the UNET trained to predict the probabilities in step 1 is shown in Table A.3.

**Table A.3:** Structure of UNET predicting per-pixel probabilities of inclusion within NFT. The left 3 columns describe the contraction layers, and the right 3 columns describe the expansion layers. The number of parameters correspond to those associated with linear filters + bias vector. Conv: 3 × 3 convolution with stride 1, MP: 2 × 2 max pool, ReLU: Rectified Linear Unit, ConvT: 2 × 2 transposed convolution with stride 2. The number of features in the expansion layers is double that of contraction layers due to skip connections that form a concatenation with the contraction layers.

| No. | Contract | Parameters | No. | Expand | Parameters |
| --- | --- | --- | --- | --- | --- |
| 1 | Conv | $8 \times 3 \times 3 \times 3 + 8$ | 18 | ConvT | $32 \times 64 \times 2 \times 2 + 32$ |
| 2 | Conv | $8 \times 8 \times 3 \times 3 + 8$ | 19 | Conv | $32 \times 64 \times 3 \times 3 + 32$ |
| 3 | MP | 0 | 20 | ReLU | 0 |
| 4 | Conv | $16 \times 8 \times 3 \times 3 + 16$ | 21 | Conv | $32 \times 32 \times 3 \times 3 + 32$ |
| 5 | ReLU | 0 | 22 | ReLU | 0 |
| 6 | Conv | $16 \times 16 \times 3 \times 3 + 16$ | 23 | ConvT | $16 \times 32 \times 2 \times 2 + 16$ |
| 7 | ReLU | 0 | 24 | Conv | $16 \times 32 \times 3 \times 3 + 16$ |
| 8 | MP | 0 | 25 | ReLU | 0 |
| 9 | Conv | $32 \times 16 \times 3 \times 3 + 32$ | 26 | Conv | $16 \times 16 \times 3 \times 3 + 16$ |
| 10 | ReLU | 0 | 27 | ReLU | 0 |
| 11 | Conv | $32 \times 32 \times 3 \times 3 + 32$ | 28 | ConvT | $8 \times 16 \times 2 \times 2 + 8$ |
| 12 | ReLU | 0 | 29 | Conv | $8 \times 16 \times 3 \times 3 + 8$ |
| 13 | MP | 0 | 30 | ReLU | 0 |
| 14 | Conv | $64 \times 32 \times 3 \times 3 + 64$ | 31 | Conv | $8 \times 8 \times 3 \times 3 + 8$ |
| 15 | ReLU | 0 | 32 | ReLU | 0 |
| 16 | Conv | $64 \times 64 \times 3 \times 3 + 64$ | 33 | Linear | $2 \times 8 + 2$ |
| 17 | ReLU | 0 |  |  |  |
|  |  |  |  | Total | 120,834 |

Accuracy of NFT detections was computed independently for each step of the algorithm. Per-pixel probabilities of tau were evaluated using 10-fold cross validation. Results for each brain sample are summarized in Tables A.4 and A.5.

**Table A.4:** 10-fold cross validation accuracy statistics for training data of brain sample 1.

| Trial | AUC | Precision | Recall | Accuracy |
| --- | --- | --- | --- | --- |
| 1 | 0.9997 | 0.0455 | 1.0000 | 0.9928 |
| 2 | 0.9983 | 0.0938 | 0.9917 | 0.9829 |
| 3 | 0.9963 | 0.1148 | 0.9846 | 0.9706 |
| 4 | 0.9984 | 0.2079 | 0.9971 | 0.9877 |
| 5 | 0.9796 | 0.3010 | 0.9242 | 0.9543 |
| 6 | 0.9915 | 0.0500 | 0.9620 | 0.9597 |
| 7 | 0.9865 | 0.2462 | 0.9031 | 0.9899 |
| 8 | 0.9239 | 0.0081 | 0.7079 | 0.9520 |
| 9 | 0.9989 | 0.1714 | 1.000 | 0.9939 |
| 10 | 0.9867 | 0.0255 | 0.9406 | 0.9454 |
| Average | 0.9860 | 0.1264 | 0.9411 | 0.9729 |

**Table A.5:**
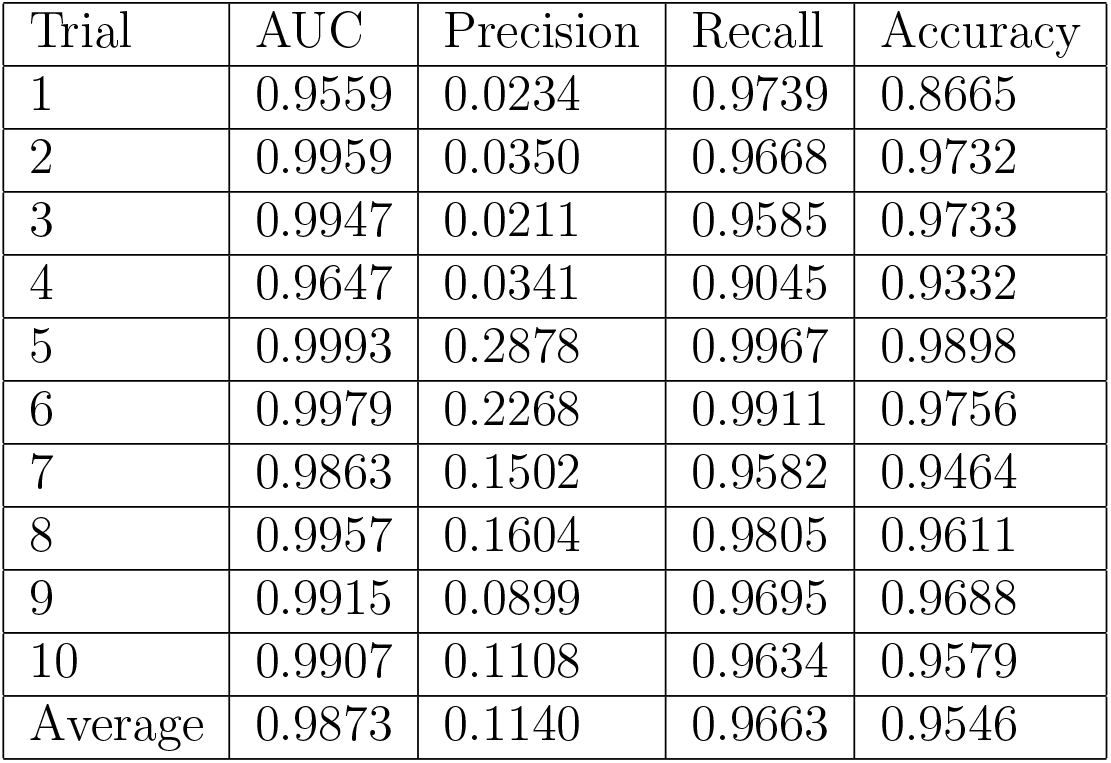
10-fold cross validation accuracy statistics for training data of brain sample 2.

| Trial | AUC | Precision | Recall | Accuracy |
| --- | --- | --- | --- | --- |
| 1 | 0.9559 | 0.0234 | 0.9739 | 0.8665 |
| 2 | 0.9959 | 0.0350 | 0.9668 | 0.9732 |
| 3 | 0.9947 | 0.0211 | 0.9585 | 0.9733 |
| 4 | 0.9647 | 0.0341 | 0.9045 | 0.9332 |
| 5 | 0.9993 | 0.2878 | 0.9967 | 0.9898 |
| 6 | 0.9979 | 0.2268 | 0.9911 | 0.9756 |
| 7 | 0.9863 | 0.1502 | 0.9582 | 0.9464 |
| 8 | 0.9957 | 0.1604 | 0.9805 | 0.9611 |
| 9 | 0.9915 | 0.0899 | 0.9695 | 0.9688 |
| 10 | 0.9907 | 0.1108 | 0.9634 | 0.9579 |
| Average | 0.9873 | 0.1140 | 0.9663 | 0.9546 |

Segmentations of probability maps into discrete NFTs were validated through direct comparison to manual annotations of NFTs in a subset of annotated image patches reserved for validation. Between 5 and 20 mm^2^ patches in the region of the ERC on 10 roughly consecutive slices of one brain were selected for annotation. Per pixel annotations were completed by a single individual over the course of 2-3 weeks. Each patch totaled approximately 2, 500, 000 pixels, yielding a total of 25 million for the 10 patches–on the order of the number of voxels for 20 whole brain MRIs at 1 mm resolution. Total counts of NFTs within each patch yielded through manual annotation versus machine detection were compared 1:1 (see Figure A.8) and demonstrated concordance both in relative trends slice-to-slice and in absolute number of NFTs per patch.

**Figure A.8:**
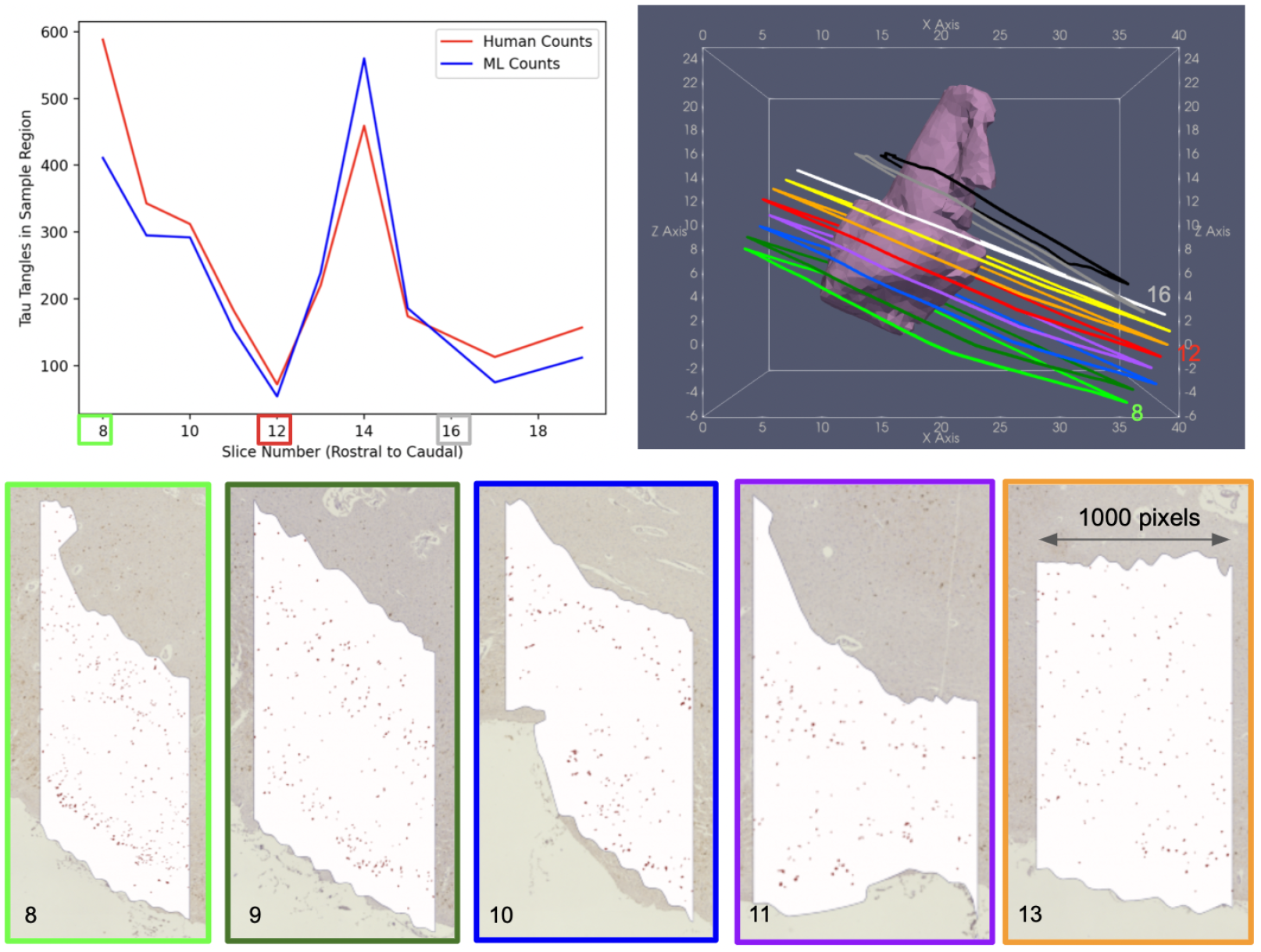
Selected histology slices with 2D segmentations (top row) ordered left to right as rostral to caudal. Corresponding MRI slices with 3D segmentations mapped to 2D via transformations *φ, ϕ*_*n*_ (second row).Counts of neurofibrillary tau tangles (NFTs) within patches of ERC computed from manual annotations (red) and machine prediction (blue) (third row, left). Outlined sections of histology from which validation set of ERC patches was taken are plotted post transformation in coordinate space of Mai atlas with 3D reconstruction of total ERC (third row, right). Example patches in ERC (white) with NFTs annotated (red) for five slices (bottom).

## Acknowledgements

Data collection, preparation, and sharing for this project was funded in part by the Alzheimer’s Disease Neuroimaging Initiative (ADNI) (National Institutes of Health Grant U01 AG024904) and DODADNI (Department of Defense award number W81XWH-12-2-0012). ADNI is funded by the National Institute on Aging, the National Institute of Biomedical Imaging and Bioengineering, and through generous contributions from the following: AbbVie, Alzheimer’s Association; Alzheimer’s DrugDiscovery Foundation; Araclon Biotech; BioClinica, Inc.; Biogen;Bristol-Myers Squibb Company; CereSpir, Inc.; Cogstate; Eisai Inc.; Elan Pharmaceuticals, Inc.; Eli Lilly and Company; EuroImmun; F.Hoffmann-La Roche Ltd and its affiliated company Genentech, Inc.;Fujirebio; GE Healthcare; IXICO Ltd.; Janssen Alzheimer Immunotherapy Research Development, LLC; Johnson Johnson Pharmaceutical Research Development LLC; Lumosity; Lundbeck;Merck Co, Inc.; Meso Scale Diagnostics, LLC; NeuroRx Research;Neurotrack Technologies; Novartis Pharmaceuticals Corporation; PfizerInc.; Piramal Imaging; Servier; Takeda Pharmaceutical Company; and Transition Therapeutics. The Canadian Institutes of Health Research provides funds to support ADNI clinical sites in Canada. Private sector contributions are facilitated by the Foundation for the National Institutes of Health (www.fnih.org). The grantee organization is the Northern California Institute for Research and Education, and the study is coordinated by the Alzheimer’s Therapeutic Research Institute at the University of Southern California. ADNI data are disseminated by the Laboratory for Neuroimaging at the University of Southern California. We thank Henry Noren for his work organizing data and running software. We thank Ophelia Yang for her work on augmenting training data for machine learning.

